# β-glucan utilization in marine *Bacteroidota* is controlled by a membrane-spanning one-component system

**DOI:** 10.64898/2026.01.10.698779

**Authors:** Daniel Bartosik, Marie-Katherin Zühlke, Norma Welsch, François Thomas, Thomas Schweder

## Abstract

One of the most prevalent and bioavailable glycans in marine systems is the β-glucan laminarin. Members of the phylum *Bacteroidota* are particularly well adapted to degrade this and other polysaccharides. Here, we describe a membrane-spanning one-component system that regulates laminarin utilization in marine *Bacteroidota*. We analyzed this β-glucan utilization regulator (BguR) type in the marine model bacterium *Formosa agariphila* KMM3901^T^. Deletion of the regulator gene abolishes growth on laminarin, whereas the wild type exhibits more than 80-fold induction of the associated genomic gene cluster, indicating the regulator’s role as a transcriptional activator for laminarin utilization. Structural predictions show that its periplasmic sensor domain resembles those of hybrid two-component systems (HTCSs), although the absence of phosphorylation domains and distinct architecture indicate a completely different, ATP-independent mode-of-action. Comparative genomics show that this regulator is widespread among *Bacteroidota*, exhibiting lineage-specific distribution patterns similar to hallmark features such as tandem SusCD-like pairs. BguR is frequently found in close proximity to β-glucan-targeting PULs in the genome, implying a defined substrate preference that extends beyond laminarin. These findings suggest a previously unrecognized regulatory mechanism for glycan sensing in marine bacteria, shedding light on an important facet of the marine carbon cycle.

**IMPORTANCE:** Although recent research has provided detailed insights into the enzymatic breakdown of algal glycans by marine bacteria, the regulatory mechanisms that govern specific carbohydrate utilization strategies remain largely unexplored. This study reveals a novel regulatory mechanism for β-glucan use by marine *Bacteroidota*. We identified a membrane-embedded one-component regulator that directly links sensing and utilization of laminarin, a major algal polysaccharide in the oceans. Unlike traditional two-component systems, the new regulator operates without phosphorelays. The regulator and its associated laminarin PULs are widely conserved across diverse marine bacteria and even linked to other β-glucan types, indicating a shared regulatory strategy for carbon acquisition from structurally related substrates. These findings advance our understanding of how key ocean microbes coordinate polysaccharide breakdown.

## INTRODUCTION

The β-1,3-glucan laminarin represents a major fraction of algal primary production and is a key source of carbon and energy for heterotrophic bacterioplankton in the ocean (1). Members of the phylum *Bacteroidota* are particularly important in utilizing this abundant marine glycan and other, more complex polysaccharides, acting as key degraders of algal-derived organic matter (2). This capacity is largely mediated by polysaccharide utilization loci (PULs), which facilitate the binding, transport and enzymatic breakdown of glycans (3). While the enzymatic degradation cascade for laminarin is well characterized (4), far less is known about the regulation of PUL expression in marine systems (5). To date, the only experimentally validated regulator of a marine PUL is a cytoplasmic GntR-type repressor controlling alginate utilization in *Zobellia galactanivorans* (6). Beyond such cytoplasmic regulators, bacteroidotal PULs are typically predicted to be controlled by hybrid two-component systems (HTCS) or extracytoplasmic function sigma factors (3, 7), yet characterization of these systems in marine representatives remains scarce.

Our genomic analysis indicated a novel transcriptional regulator that controls laminarin utilization. Comparative genomics reveals widespread homologs of this regulator in global marine metagenomes. Our integrated proteogenomic, physiological, and computational structural biological analyses elucidate the *in situ* function of this regulatory system and its ecological relevance.

## RESULTS AND DISCUSSION

### BguR controls laminarin utilization

The bacteroidotal model strain *Formosa agariphila* KMM3901^T^ degrades branched laminarin via dedicated PUL encoding glycoside hydrolases of the GH16_3, GH17, GH30_1, GH149 and GH158 families (Fig. 1A), all of which have been reported to be involved in laminarin degradation (4, 8–10). Immediately upstream of the SusCD-like transporter pair, we identified a gene encoding a putative membrane-embedded transcriptional regulator, currently annotated as a ‘two-component system response regulator, LuxR family’ (UniProt ID T2KMC4) (11), here designated BguR (β-glucan utilization regulator).

**Fig 1.**
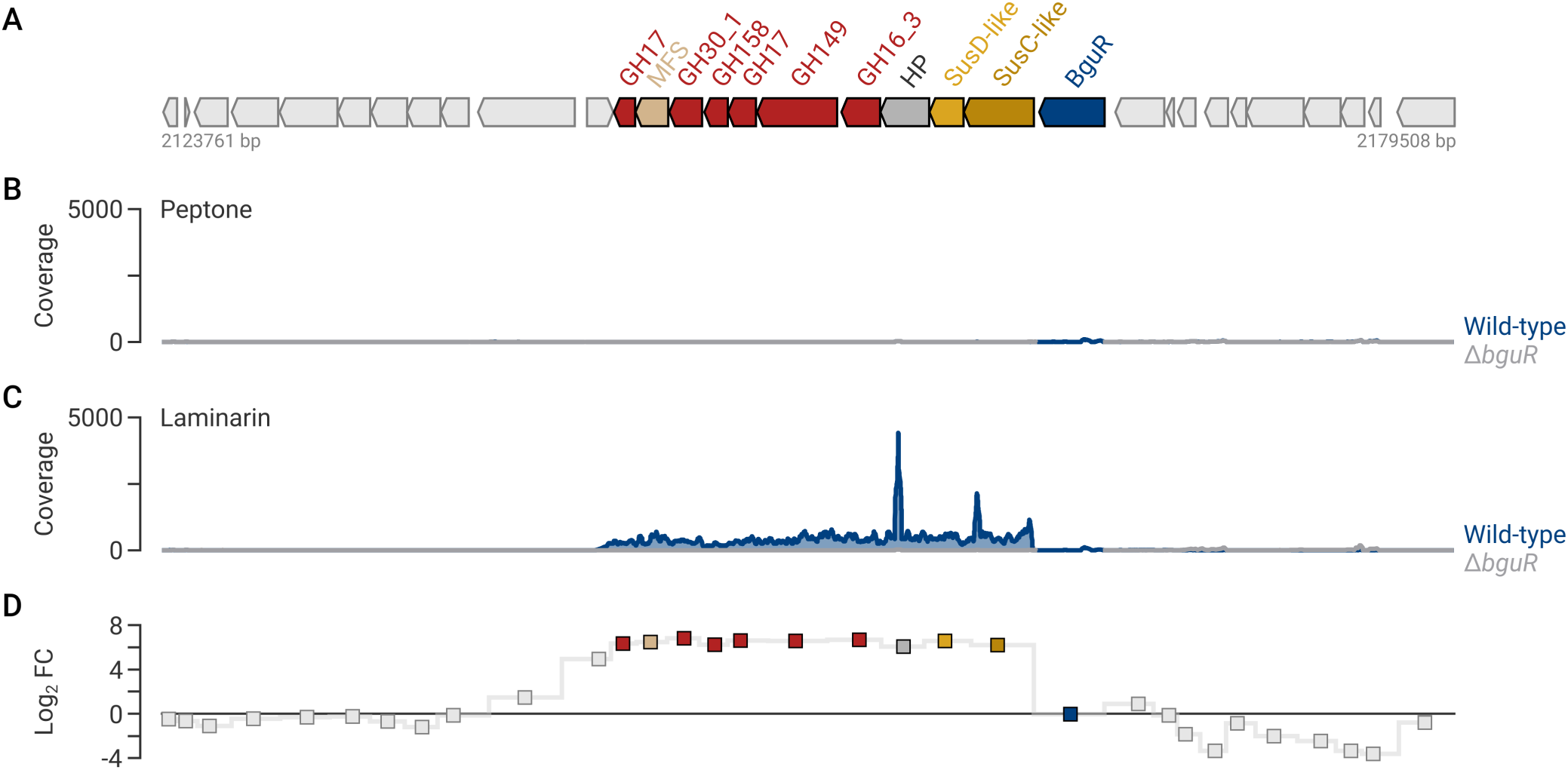
(A) Organization of the *F. agariphila* KMM3901^T^ laminarin PUL. PUL-encoded proteins are colored according to function: CAZymes (red), transporters (yellow shades), and the regulator BguR (blue). Ten genes upstream and downstream of the PUL are shown for genomic context. HP: hypothetical **(B)** Read coverage per nucleotide across the laminarin PUL and neighboring genes for the wild-type and Δ*bguR* grown on peptone (n = 3). **(C)** Read coverage per nucleotide across the laminarin PUL and neighboring genes for the wild-type and Δ*bguR* grown on laminarin (n = 3). **(D)** DESeq2-based interaction analysis of laminarin PUL gene expression (n = 3) 3 h after a substrate shift from peptone to laminarin, testing for a significant interaction between genotype (wild-type vs. Δ*bguR*) and carbon source (laminarin vs. peptone).

To assess the role of BguR in laminarin utilization, growth of the *F. agariphila* KMM3901^T^ wild-type and the Δ*bguR* mutant was compared on glucose, alginate, and laminarin derived from *Laminaria digitata* and *Eisenia bicyclis*. On glucose and alginate, both strains showed nearly identical growth kinetics (Fig. 2A,B), reaching the stationary phase after approximately 24 h and achieving comparable maximum optical densities (glucose: WT OD₆₀₀ ≈ 2.1, Δ*bguR* ≈ 1.9; alginate: WT ≈ 1.16, Δ*bguR* ≈ 1.07), indicating that loss of *bguR* does not affect general growth behavior or utilization of these carbon sources. In contrast, the Δ*bguR* mutant failed to grow on laminarin from either algal source (Fig. 2C,D). Whereas the wild-type reached OD₆₀₀ values of approximately 1.72 on laminarin from *L. digitata* and 1.21 from *E. bicyclis*, the Δ*bguR* mutant showed only a minor increase in optical density during the first 24–32 h (OD₆₀₀ ≤ 0.20). Thereafter, cell densities remained constant or slightly decreased, suggesting that the initial increase in the optical density resulted from residual carbon carried over from the pre-culture rather than laminarin utilization. The wild-type showed substrate-dependent growth characteristics, displaying a higher growth rate (μ = 0.15 h⁻¹ versus 0.096 h⁻¹), and a higher final cell density on *L. digitata* laminarin (OD₆₀₀ ≈ 1.7 after 24 h) than on *E. bicyclis* laminarin (OD₆₀₀ ≈ 1.2 after 32 h), consistent with the higher degree of β-1,6 branching reported for *E. bicyclis* laminarin (13), which likely hampers enzymatic degradation. Taken together, these results demonstrate that BguR is indispensable for growth on laminarin in *F. agariphila* KMM3901^T^.

**Fig 2.**
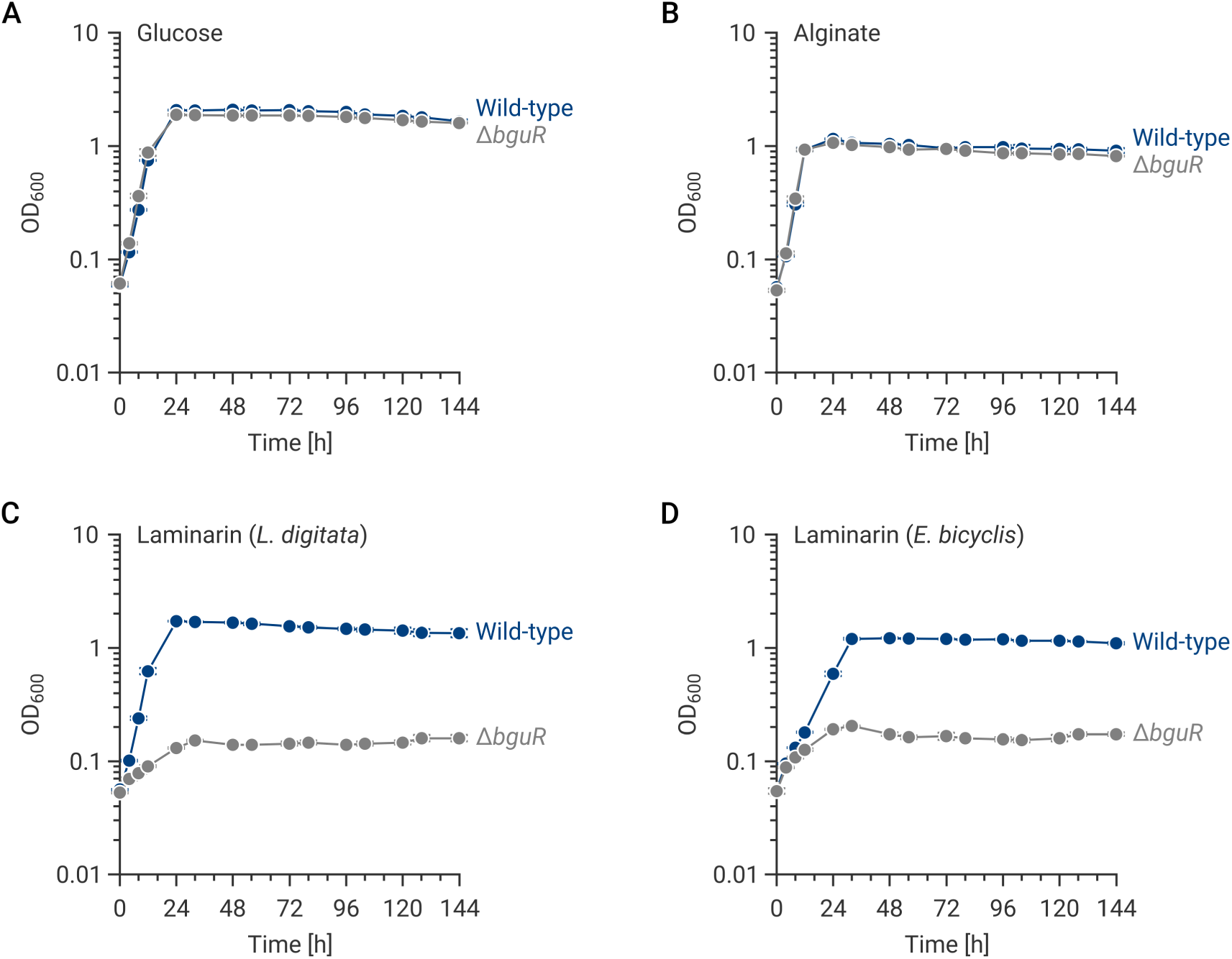
Growth curves of *F. agariphila* KMM3901^T^ wild-type and Δ*bguR* strains cultivated on **(A)** glucose, **(B)** alginate, **(C)** laminarin from *L. digitata* and **(D)** laminarin from *E. bicyclis* as sole carbon sources (synthetic seawater medium (MPM) supplemented with 0.2% (w/v) substrate). Cultures were incubated at 25°C and 250 rpm for 144 h. Data represent mean ± SD of three biological replicates.

Because growth phenotypes alone cannot distinguish whether BguR functions as a transcriptional activator or a repressor, we next examined the transcriptional expression of laminarin PUL genes in the wild-type and Δ*bguR* strains during growth on laminarin as the sole carbon source following pre-cultivation in a peptone-containing medium. Read coverage showed virtually no transcription across the genes of the PUL during growth on peptone in either strain (Fig. 1B). In contrast, cultivation on laminarin resulted in strong gene transcription across the entire PUL in the wild-type, whereas transcript levels remained close to background levels in the Δ*bguR* mutant (Fig. 1C). Differential expression analysis using DESeq2 (12) confirmed the coverage-based observations, showing strong induction of the laminarin PUL in the wild-type relative to the Δ*bguR* mutant. PUL specific genes exhibited log₂ fold changes between 6.1 and 6.8, corresponding to approximately 70- to 110-fold higher transcript abundance (Fig. 1D). Notably, the gene encoding a hypothetical protein (NCBI locus_tag BN863_RS18030) contains so far unclassified carbohydrate-binding modules which are distantly related to the CBM103 family (13), suggesting a role in laminarin binding.

### BguR is a membrane-spanning one-component regulator

BguR is a 940-aa membrane-spanning protein that contains a single transmembrane domain separating a putative periplasmic sensor from a cytoplasmic DNA-binding region (Fig. 3A). Structural prediction revealed that the sensor adopts an architecture composed of two β-propeller regions, closely resembling the periplasmic sensor domains of hybrid two-component system (HTCS) regulators, known for PUL-regulation in gut-associated *Bacteroidota* (10,11). Structural superposition with the periplasmic sensor domain of an HTCS regulator from *Bacteroides thetaiotaomicron* confirmed conservation of this characteristic tandem β-propeller architecture (Fig. 3B) despite the poor overall structural similarity (RMSD ≈ 13.8 Å). However, the membrane-proximal Y_Y_Y-repeat (PF07495), a hallmark of HTCS sensory domains, is highly similar (RMSD ≈ 1.6 Å; Fig. 3B), indicating that the structural core of the sensor is retained. In contrast to canonical HTCS regulators, the cytoplasmic region of BguR lacks the conserved HisKA (PF00512), HATPase_c (PF02518), and response regulator (PF00072) domains required for ATP-dependent phosphorelay (7). Instead, the periplasmic sensor is directly connected via an α-helical linker to a GerE-like helix-turn-helix DNA-binding domain (PF00196) (Fig. 3D). The complete absence of phosphotransfer modules indicates that signal perception is not relayed through a classical phosphorylation cascade. Instead, periplasmic substrate sensing is likely coupled directly to transcriptional regulation, classifying BguR as a membrane-embedded one-component system (OCS) (12). XSTREME analysis identified a statistically significant sequence motif (E-value = 2.4e-85) representing a putative BguR binding site upstream of the first structural gene in BguR-associated β-glucan PULs (Fig. 3C).

**Fig 3.**
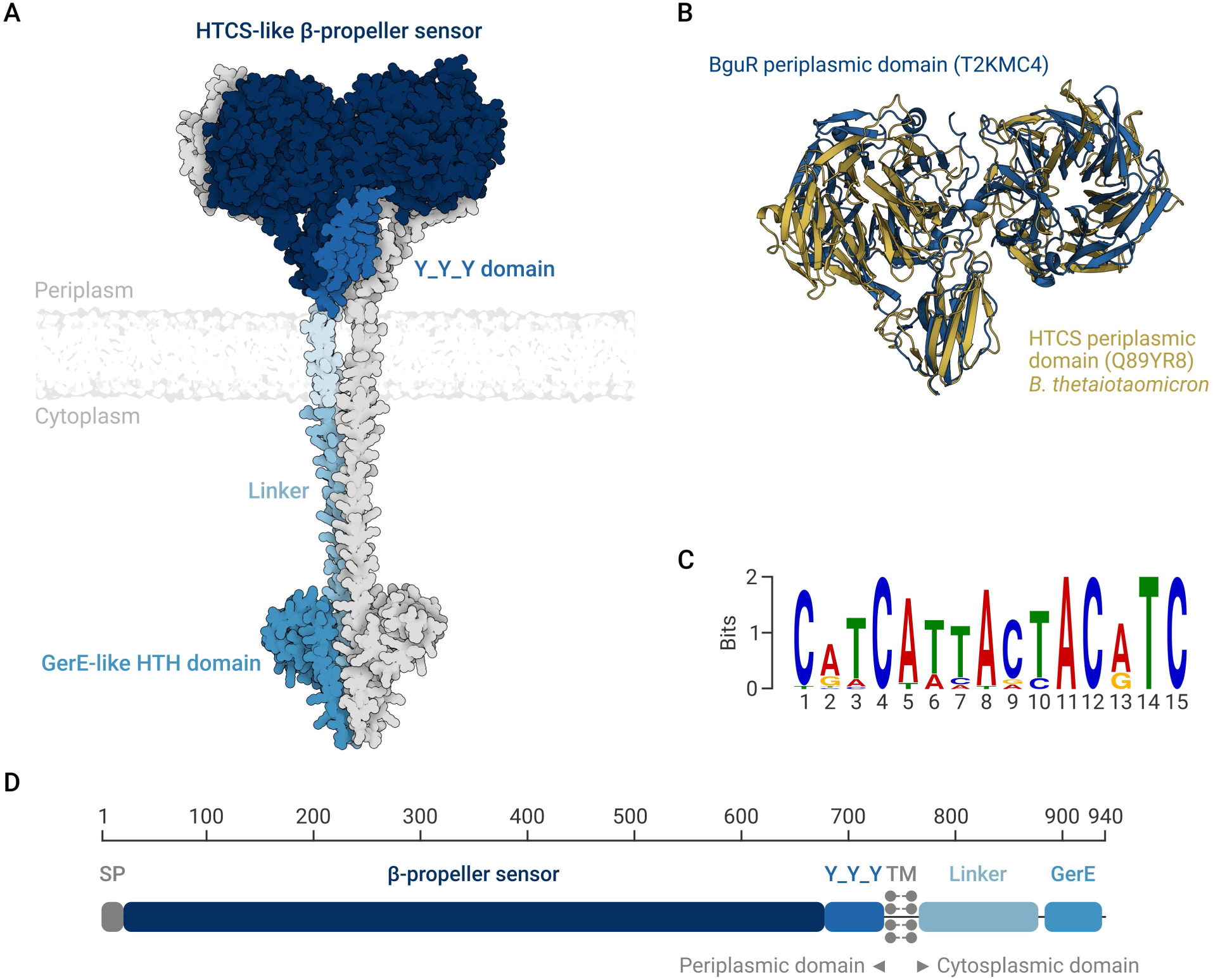
The membrane-spanning OCS regulator BguR from *F. agariphila* KMM3901^T^. **(A)** AlphaFold3-predicted structure of the BguR dimer. Structurally distinguishable regions are highlighted in shades of blue; the second monomer is shown in grey. **(B)** Superimposition of the AlphaFold3-predicted structure of the HTCS sensor domain from *B. thetaiotaomicron* (gold) with the BguR sensor domain from *F. agariphila* (blue). UniProt accessions are given in parentheses. (**C**) Conserved putative BguR-binding motif identified by XSTREME (E-value = 2.4e-85). The motif was detected in intergenic regions between *bguR* homologs and the first structural gene of associated β-glucan PULs. Letter heights are proportional to nucleotide conservation. **(D)** Schematic representation of the domain architecture of BguR. The protein contains an N-terminal signal peptide (SP), a β-propeller sensor domain, a Y_Y_Y domain, a transmembrane (TM) domain separating the periplasmic and cytoplasmic regions, a linker, and a C-terminal GerE domain. Amino acid positions are shown above the schematic.

A comparable regulatory architecture has so far only been described for SusR, the archetypal regulator of the starch utilization system in gut *Bacteroidota* (3). Like BguR, SusR lacks phosphorelay domains and represents a membrane-associated OCS, although its molecular mechanism remains unresolved (Fig. S1). While SusR is associated with sensing starch, an α-linked glucose polymer, BguR regulates the utilization of β-linked glucans such as laminarin, suggesting that membrane-associated one-component systems can be employed to control the utilization of chemically distinct, yet simple glucose-based polysaccharides.

Compared with canonical HTCS regulators, which rely on ATP-dependent autophosphorylation and phosphotransfer to connect periplasmic sensing with transcriptional output, the simplified architecture of BguR likely enables direct signal transmission without an ATP-dependent phosphorelay. This streamlined signaling mechanism may reduce both energetic costs and the delay associated with multistep phosphotransfer, potentially allowing a rapid transcriptional response to fluctuating laminarin availability during episodic algal blooms in marine environments.

### BguR homologs diverge according to β-glucan specificity

Given the central role of BguR in laminarin utilization in *F. agariphila* KMM3901^T^, we next investigated the distribution of BguR homologs across marine *Bacteroidota*. Screening of 53 genomes (15) identified BguR homologs exclusively within β-glucan-targeting PULs. Besides the expected association with laminarin utilization loci encoding GH3, GH5_46, GH16, GH17, GH30, GH149 and GH158 family enzymes (16), BguR homologs were also consistently associated with GH144-containing PULs involved in β-1,2-glucan degradation (17,18) (Fig. 4). No associations with PULs targeting other polysaccharides were detected. Phylogenetic analysis resolved the BguR homologs into two major clades corresponding to substrate specificity, separating regulators associated with laminarin PULs from those linked to β-1,2-glucan utilization loci. Across all homologs, the mean pairwise amino acid identity is 47.9% (median 45.8%). Within the laminarin-associated clade, homologs showed a comparable level of conservation (mean 46.7%, median 45.3%), whereas β-1,2-glucan-associated BguR proteins were considerably more similar to one another (mean 58.9%, median 54.4%). Kappelmann *et al*. (2019) suggested a separation of three different laminarin PUL types (A, B and C) according to the CAZyme composition in marine *Bacteroidota* (14). Despite substantial variation in CAZyme composition, laminarin-associated BguR homologs showed little segregation according to type A and B laminarin specific PUL architecture, with only type C laminarin PULs (encoding GH5_46, GH16_3, GH30_1 and GH30_3) forming a partially distinct subgroup. The strict association of BguR homologs to β-glucan utilization loci, together with their separation into substrate-specific clades rather than PUL architectures, suggests that BguR represents a conserved regulatory module that diversified according to β-glucan substrate specificity while remaining largely independent of the enzymatic composition of individual PULs.

**Fig 4.**
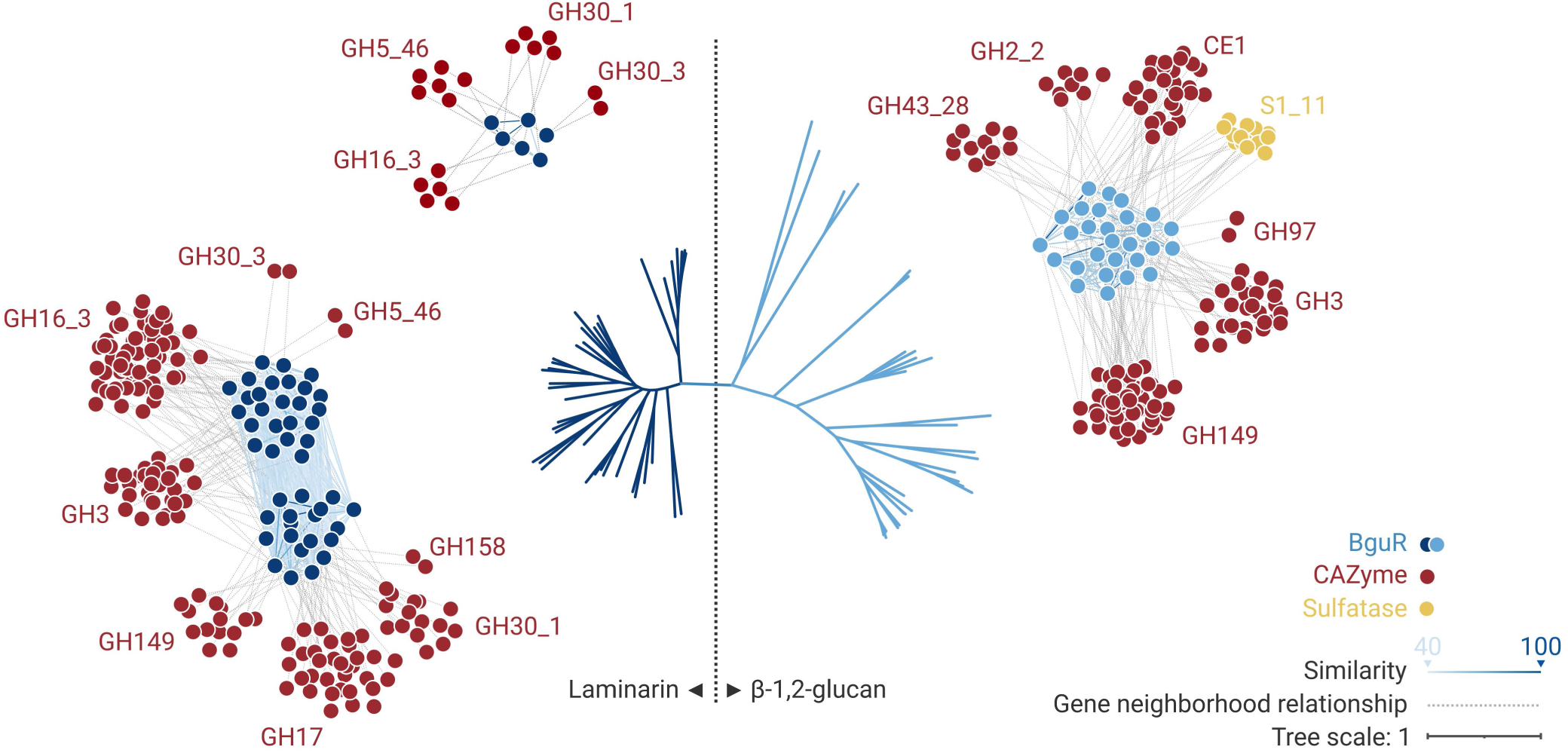
Phylogenetic relationships and network analysis of BguR-associated PULs in 54 marine *Flavobacteriia*. The phylogenetic tree is based on BguR amino acid sequences. Dashed lines represent co-occurrence of BguR-associated CAZymes and sulfatases within PULs, whereas colored lines indicate sequence identity between BguR proteins.

To determine whether this conserved association translates into ecological relevance, we queried the Ocean Microbial Reference Gene Catalog (15, 16). Homologs of BguR were detected across marine metagenomes from all major oceanic regions, spanning polar, temperate, and tropical waters as well as both coastal and open-ocean environments (Fig. 5). Relative abundances ranged from 6.28 × 10⁻¹⁰ to 1.39 × 10⁻⁴, indicating that although local abundances varied substantially, BguR is globally distributed throughout the world’s oceans. Taxonomic profiling further showed that its occurrence is almost as tightly restricted to *Bacteroidota* (94.1 - 98.6%) (Fig. S2) as canonical SusCD-like proteins (94.4 - 98.6%) (3, 17), mirroring their phyletic distribution and pointing to a conserved, phylum-level regulatory component underlying β-glucan utilization.

**Fig 5.**
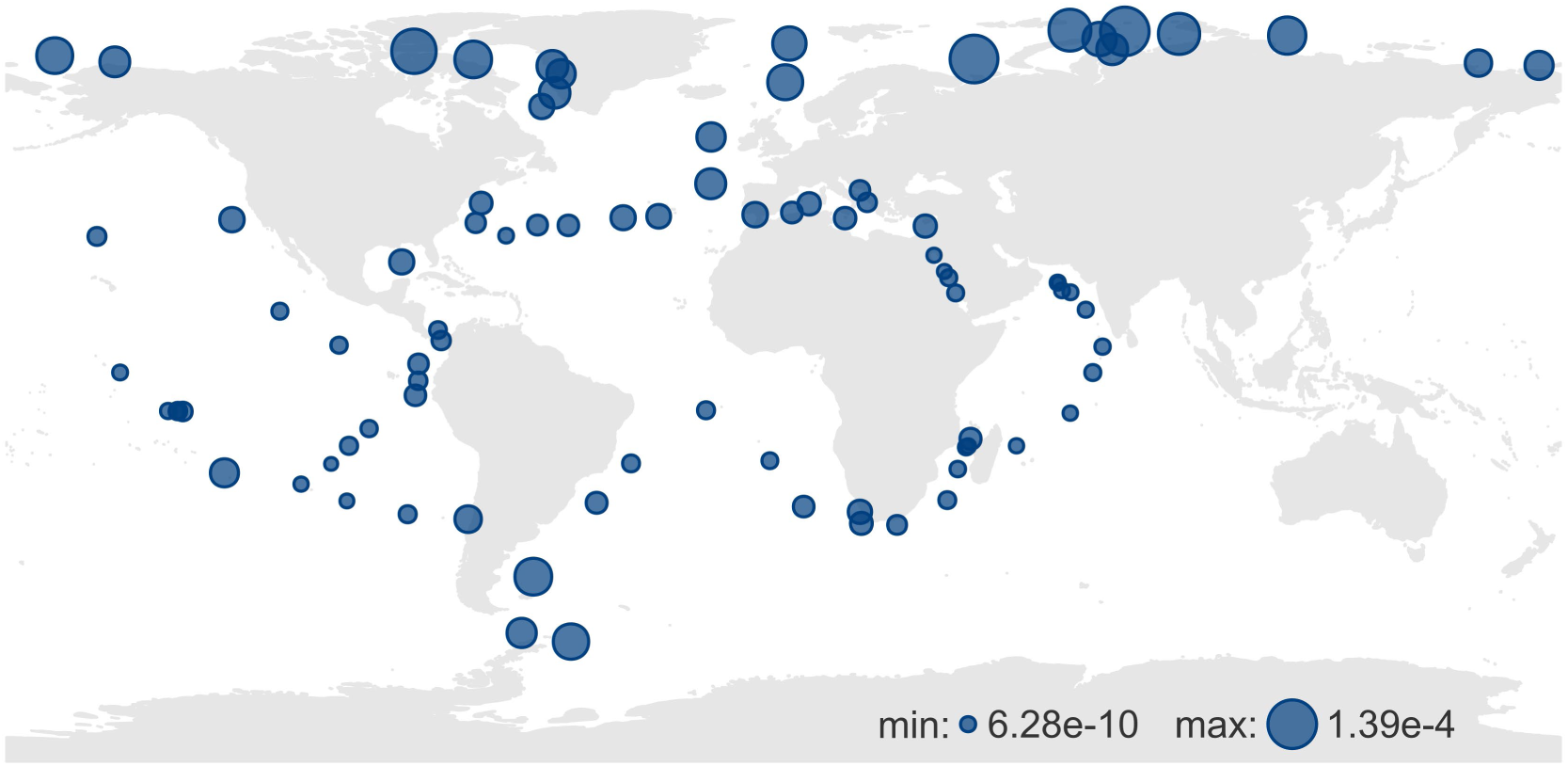
Global distribution of BguR homologs based on the Ocean Gene Atlas v2.0, using the Tara Oceans Microbiome Reference Catalog v2+ MetaG Arctic Inside (prokaryotes) dataset. Bubble sizes correlate with the percentage of mapped reads.

The results obtained suggest BguR as an ATP-independent, membrane-embedded one-component regulator that directly couples periplasmic β-glucan sensing to transcriptional activation of a laminarin specific PUL expression. This strategy is distinct from canonical HTCS-based transcriptional control. Comparative genomics shows a consistent association of BguR homologs not only with laminarin PULs but also with loci putatively targeting β-1,2-glucans. The widespread occurrence of BguR homologs in global marine metagenomes indicates that this regulatory architecture is conserved for controlling β-glucan utilization in *Bacteroidota* across marine environments.

## MATERIALS AND METHODS

### Deletion of the BguR encoding gene

We used the BguR encoding gene (locus tag BN863_18790) of *F. agariphila* KMM 3901^T^ as model to investigate the function of this putative bacteroidotal β-glucan regulator. Construction of the appropriate *bguR* deletion vector was achieved by Gibson assembly. To this purpose, PCR reactions of both BguR homology regions HomA (upstream of the start-codon) and HomB (downstream of the stop-codon) were carried out with oligonucleotides fw front BguR/rev front BguR (HomA) and fw back BguR/rev back BguR (HomB) (Table S1) using genomic DNA of *F. agariphila* KMM3901T as the template. The vector backbone of pYT313 (18) was amplified in two PCRs with oligonucleotides fw pYT313_1/rev pYT313_3 and fw pYT313_2/rev pYT313_4. The PCR products were gel-purified using the Qiagen gel extraction kit and all four DNA fragments were then assembled in a vector:insert ratio of 1:2. 3 µL of the reaction were used for transformation of *E. coli* DH10B yielding pYT313-HomAB-BguR. Sequence identity of the deletion vector was verified and the plasmid was subsequently integrated into *E. coli* S17λpir by electroporation.

In principle, conjugation procedure was carried out as described for *Z. galactanivorans* DsiJ^T^ (18). Due to the intended genomic modification of *F. agariphila*, adaptions/changes in media composition and cultivation conditions were necessary. In brief, the plasmid-harbouring donor strain *E. coli* S17λpir was grown overnight in Luria Broth (LB) at 37°C, whereas the *F. agariphila* recipient strain was grown overnight in Marine Broth (MB) at 25°C. For both strains, cells were harvested by centrifugation and washed in the respective cultivation medium. Donor- and recipient strains were re-suspended in marine conjugation medium prior to filter mating on marine conjugation agar. Plates were incubated at 25°C overnight. The next day, cells were scraped off from the filter and diluted in MB before plating them on erythromycin containing MB agar. Plates were incubated for 4-5 d at 25°C. Due to a single cross-over event, erythromycin resistant colonies should contain the deletion vector pYT313-HomAB-BguR within the genome and these colonies were then streaked again on MB-agar with erythromycin in order to obtain single colonies. One single colony was cultivated in MB medium without erythromycin and cells were plated on MB agar with 5% sucrose. In order to obtain plasmid-free cells, colonies grown on sucrose were streaked out again in the presence of sucrose for isolation of single cell colonies. Colony PCRs with oligonucleotides fw BguR del and rev BguR del were carried out to screen these single colonies for the correct deletion. Selected putative deletion mutants were then grown in MB, genomic DNA was isolated and deletion of the BguR coding sequence was verified by PCR using purified gDNA as the template. Additionally, the PCR products were sequenced to confirm the correct genotype.

### Cultivation of *F. agariphila* strains with different carbon sources

*F. agariphila* wild-type and BguR deletion strains were cultivated in presence of different carbon sources: glucose, alginate (both serving as the control), laminarin from *Laminaria digitata* and laminarin from *Eisenia bicyclis*. Synthetic seawater medium (MPM) (19), supplemented with the respective substrate (in a final concentration of 0.2%), was used as the growth medium. Both strains were cultivated at 25°C and 250 rpm for 6 d (144 h).

### Cultivation of *F. agariphila* strains for transcriptome analysis

To gain insights into differences in the expression level between wild-type and Δ*bguR* strain, cells were initially cultivated in MPM supplemented with 0.2% peptone before they were transferred to MPM with 0.2% laminarin (from *L. digitata*). Samples for RNA isolation were harvested immediately before and 3 h after substrate transition normalized on cell count. In detail, triplicates of both strains were grown to an OD_600nm_ of 0.6 at 25°C and 250 rpm in MPM-peptone. Cells were collected by centrifugation of the entire culture volume at 4°C and 5,000 x g for 30 min. The supernatant was discarded, the cell pellets were washed in substrate-free MPM basal medium and centrifuged. Subsequently, the pellets were re-suspended in 750 µL of MPM without substrates and the entire amount of cells was transferred into 50 mL of fresh MPM supplemented with 0.2% laminarin. Growth was continued for another 3 h before cells were harvested for RNA isolation. Cell pellets were mixed with RNA later (Sigma/Merck), shock-freezed in liquid nitrogen and stored at −80°C until RNA isolation.

### RNA isolation

Prior to mRNA sequencing, total RNA was isolated by a combined approach of cell lysis using Tri Reagent (Sigma/Merck) and the Monarch® Total RNA Miniprep Kit (NEB). Both procedures were carried out as recommended by the manufactureŕs instructions. Briefly, the thawed cell pellets were centrifuged at 4°C and 5,000 x g for 5 min and the pellets were homogenized in 1 mL of Tri Reagent and incubated for 5 min at RT. Subsequently, 200 µL of chloroform were added and samples were shaken vigorously for 15 sec. After an additional incubation step (15 min, RT), samples were centrifuged at 4°C and 12,000 x g for 15 min. The upper aqueous phase was then subjected to RNA isolation using the Total RNA Miniprep Kit. After the purification procedure, the RNA was eluted in 50 µL of RNase-free water and the RNA concentration was determined using the Nanodrop ND1000 device.

### Transcriptome analysis

For prokaryotic mRNA sequencing, 20 µL of the RNA samples were sent to Biomarker Technologies (BMK) GmbH (Münster, Germany). All necessary steps prior to the sequencing (quality control, rRNA depletion) were carried out at BMK. Sequencing results were provided as raw data and were then further evaluated (Supplementary Data Set 1).

Raw sequencing reads of three biological replicates per sample were quality checked with FastQC v0.12.1 (20). Adapter sequences and low-quality bases were trimmed with Trim Galore v0.6.10 (21) invoking Cutadapt v5.1 (22), using a Quality Phred score cutoff of 20, adapter sequence ‘AGATCGGAAGAGC’, a maximum trimming error rate of 0.1 and a minimum required sequence length of 20 bp. rRNAs and tmRNAs were removed using SortmeRNA v4.3.7 (23) with default settings and --ref smr_v4.3_default_db.fasta and --ref RF00023.fa. Remaining paired-end reads were mapped on *F. agariphila* KMM3901^T^ reference genome ‘GCF_000723205.1’ (19) from NCBI (24) using Bowtie2 v2.5.4 (25) with default settings. Read coverage tracks were generated from sorted BAM files using bamCoverage from the deepTools2 package and exported in BigWig format for visualization (26). Finally, paired-end reads per coding DNA sequence (CDS) were quantified with featureCounts v2.0.0 (27). Differential gene expression (wild-type vs *ΔbguR*; peptone- vs-laminarin-containing medium) was analyzed with the DESeq2 package (12).

### PUL and CAZyme prediction

CAZymes in bacteroidotal genomes (14) were predicted using hmmscan (HMMER 3.3.2) (28) against dbCAN-HMMdb-V14 and dbCAN-sub as well as diamond blastp v2.1.1.155 (29) using flags --evalue 1E-20 --id 30 --query-cover 40 against CAZyDB.07242025, provided by dbCAN3 (30). Results were further parsed using the hmmscan-parser.sh script with an e-value cutoff of 1E-5 and a minimum coverage of 0.3. CAZyme family assignments predicted by at least two tools were kept for further analysis. PULs were predicted as described previously (31) using a five-gene sliding window and were assigned to laminarin PUL-types according to Krüger *et al*. (17). Non-β-glucan targeting CAZyme annotations within the five-gene window of laminarin/GH144-PULs were excluded from analysis.

### BguR annotation and HMM creation

The domain architecture of *F. agariphila* KMM3901^T^ BguR (NCBI locus_tag: BN863_RS09390) was determined using hmmscan against Pfam-A database release 37.4 (32). Results were filtered using the hmmscan-parser.sh script. Transmembrane domains (TMs) were predicted with (Poly)Phobius (33). To generate an in-house Hidden Markov Model (HMM) profile for BguR screening, CDS from 53 *Flavobacteriia* genomes (14) were annotated using hmmscan and phobius as described above. BguR homologs containing a single transmembrane domain as well as Pfam profiles ‘GerE’ (PF00196.25) and ‘Y_Y_Y’ (PF07495.19) (n=53) were aligned using the m-coffee webserver (34) with default parameters. A representative set of sequences was extracted from the alignment (n=31) with hhfilter (35) as implemented in the MPI Bioinformatics Toolkit (36) applying a maximum sequence identity of 90%, a minimum sequence identity of 40% and a minimal coverage of 35%. The resulting alignment was trimmed with trimAl v1.5.0 with the ‘-automated1’ option, and a seed BguR HMM profile was constructed using hmmbuild. Using hmmsearch, 26 additional BguR homologs were identified and incorporated to refine the seed HMM profile following the same procedure as described above.

### Screening for BguR homologs in public databases

The final BguR HMM was used to screen the UniProtKB database (r2025_4) (37), the proGenomes representative protein database v3 (38) and the OM-RGC_v2 catalog (39) using hmmsearch. Hits with a minimum coverage of 70% and a length between 850-1,050 aa were retained for further analysis. To exclude false-positive hits, remaining sequences were validated using hmmscan against Pfam-A r37.4. Sequences containing cytoplasmic domains of (hybrid) two-component systems (His_kinase (PF06580.19) or HATPase_c (PF02518.31)) were excluded from subsequent analysis. To compare taxonomic specificity of BguR assignments to SusC-like (TIGR04056) and SusD-like (PF07980.15, PF12741.11, PF12771.11 or PF14322.10) proteins, all databases were screened for these HMM profiles with hmmsearch using the profile-specific gathering threshold (Supplementary Data Set 2).

### Phylogenetic tree of BguR homologs and co-occurrence similarity network

BguR homologs (Supplementary Data Set 2) were aligned using the M-Coffee web server (40) with default parameters, and maximum-likelihood phylogeny was estimated from the resulting multiple sequence alignment using the PhyML web server (41). Pairwise sequence identities between BguR proteins were determined by global alignment using EMBOSS Needle v6.6.0.0 (42, 43), and used to construct a sequence similarity network (Supplementary Data Set 3). To relate regulator divergence to PUL composition, this network was combined with a co-occurrence network of CAZyme families encoded by the corresponding BguR-associated β-glucan PULs.

### BguR binding-site prediction

Putative BguR binding sites were identified by analyzing the intergenic regions separating *bguR* homologs from the first structural gene of the associated β-glucan PULs. Sequences were screened for enriched motifs using XSTREME v5.5.9 (MEME Suite webserver) (44, 45) with default parameters. The highest-scoring conserved MEME motif was subsequently examined as a candidate BguR binding site.

### Data visualization

All shown protein structures were predicted using the AlphaFold 3 webserver (46) and visualized using PyMOL Molecular Graphics System v3.1.6.1 (Schrödinger, LLC.). Figure 1A was created using ‘Illustrate’ (47). Visualization of global BguR distribution in the Ocean Gene Atlas v2 (15) was performed in R v4.2.0 (48) using the packages ggplot2 (49), sf (50), rnaturalearth (51), dplyr (52), and ggrepel (53). Co-occurrence- and similarity network visualization was performed using Cytoscape v3.10.4 (54). The phylogenetic tree was visualized in iTOL v7.6 (55).

## Supplemental Material and Data availability

The raw RNA sequencing data generated in this study are available from the NCBI Sequence Read Archive (SRA) under BioProject accession number PRJNA1473389.

All supplementary materials, including Supplementary Figures, Supplementary Tables, and Supplementary Data files, are provided as part of the published article.

ASM does not own the copyrights to Supplemental Material that may be linked to, or accessed through, an article. The authors have granted ASM a non-exclusive, world-wide license to publish the Supplemental Material files. Please contact the corresponding author directly for reuse.

## Acknowledgements

We thank the Deutsche Forschungsgemeinschaft (DFG) for funding this study in the frame of the research unit FOR 2406 (POMPU) and the transregional CRC 420 “Carbon sequestration at Å resolution - CONCENTRATE” through grants received by Thomas Schweder (SCHW 595/10-3, Project-ID 542264307) and Marie-Katherin Zühlke (Project-ID 542264307).

## Author contributions

DB and TS designed the study supported by FT. MZ visualized protein structures/superimpositions, supported lab experiments and refined the manuscript. NW conducted mutagenesis, cultivation experiments and RNA isolation. DB performed the bioinformatic analyses including transcriptomics and wrote the manuscript with input from TS. All authors reviewed and approved the manuscript.

## Competing interest statement

The authors declare no competing interests.

## Ethical approval

Not applicable.

## Notes

### Competing Interest Statement

The authors have declared no competing interest.

### Summary of Updates

We have changed the paper's format from a Short Communication to the standard Research Paper format, which includes an introduction, results and discussion, and materials and methods section. In addition, we have expanded the two previously condensed figures into five figures to present our results in a more readable manner.

