## Supplementary Information for "β-glucan utilization in marine *Bacteroidota* is controlled by a membrane-spanning one-component system"

**The Supporting Information file contains**

Supplementary Table S1

Supplementary Figures S1 & S2

**Supplementary Data Sets 1 - 3 are provided as separate files:**

Supplementary Data Set 1: DeSeq2 data

Supplementary Data Set 2: Detailed information on BguR homologs including sequence

Supplementary Data Set 3: Alignment and co-occurrence data used in Figure 2A

**Table S1** Primer sequences used in this study

| ***Primer*** | ***Sequence 5´→3´*** |
|---|---|
| *fw front BguR* | *CCCGAAGCAGGGTTATGCAGCGGAAAAATTCGGGGGATCCCGATTTATTGATAGTTGATGAAGCTCATCGTTTAAGAC* |
| *rev front BguR* | *GATTGTATTGGTTATCTAGAGGGTTAGGTATTAGTTTGTTTAAAATGTAATATAG* |
| *fw back BguR* | *CTAATACCTAACCCTCTAGATAACCAATACAATCTACAAAATCACCCATACATTAC* |
| *rev back BguR* | *CTATGACCATGATTACGCCAAGCTTGCATGCCTGCAGGTCGACCGATTTTGCTTCACTTAATCCGTTTGCTAAAGAAATATC* |
| *fw pYT313_1* | *GTCGACCTGCAGGCATGCAAGCTTGGCGTAATCATGGTCATAGCTGTTTCCTGTGTGAAATTGTTATCCGCTCAC* |
| *fw pYT313_2* | *GTGCAATGTTGAAGATTAGTAATTCTATTCAACATTTGTGCTAAAAGTCGGCTCCATCGCCAATTTGCCAGCCGTTATGC* |
| *fw pYT313_3* | *GGCAAATTGGCGATGGAGCCGACTTTTAGCACAAATGTTGAATAGAATTACTAATCTTCAACATTGCACAAAAGTTC* |
| *fw pYT313_4* | *CATCAACTATCAATAAATCGGGATCCCCCGAATTTTTCCGCTGCATAACCCTGCTTCGGGGTCATTATAGCG* |
| *fw BguR del* | *CCAAGATGATTTAATTGTACTCTATAC* |
| *rev BguR del* | *CCAGTTTCATCTTGAACAGTTCCTGACAC* |

### Supplementary Figures

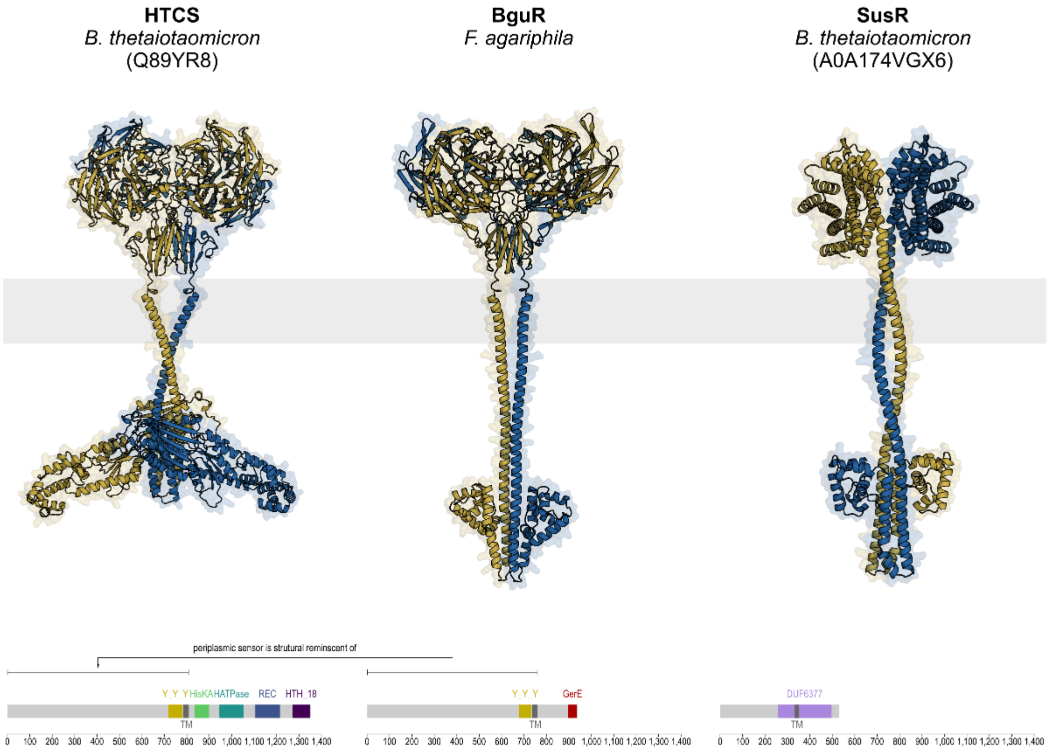


**Fig S1** AlphaFold3-predicted structures of HTCS (*B. thetaiotaomicron*), BguR (*F. agariphila*), and SusR (*B. thetaiotaomicron*). The corresponding domain architectures (Pfam-A v37.4) and protein lengths are shown below each structure.

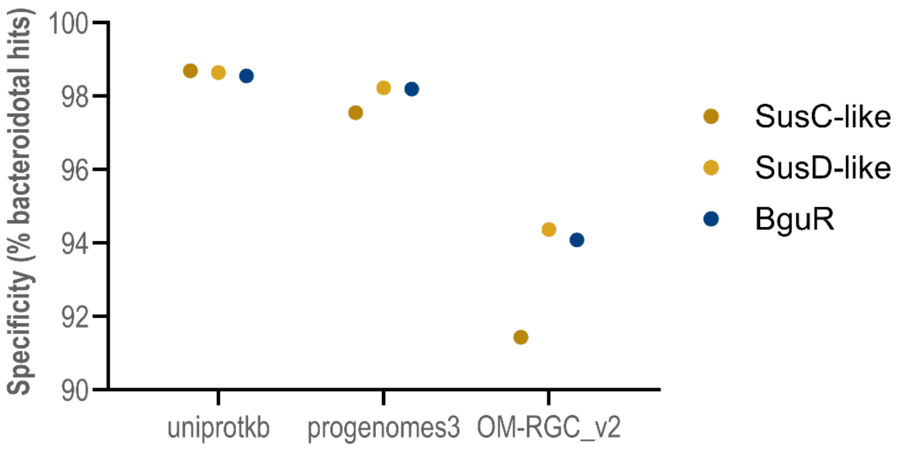


**Fig S2** Taxonomic specificity of the *Bacteroidota* marker proteins SusC-like and SusD-like compared to BguR homologs, shown as the percentage of hits from different databases assigned to the phylum *Bacteroidota*.
